# Detecting and Isolating Mass Balance Errors in Reaction Based Models in Systems Biology

**DOI:** 10.1101/816876

**Authors:** Woosub Shin, Joseph L. Hellerstein

## Abstract

**Motivation:** The growing complexity of reaction-based models necessitates early detection and resolution of model errors. This paper addresses mass balance errors, discrepancies between the mass of reactants and products in reaction specifications. One approach to detection is atomic mass analysis, which uses meta-data to expose atomic formulas of chemical species. Atomic mass analysis isolates errors to individual reactions. However, this approach burdens modelers with expressing model meta-data and writing excessively detailed reactions that include implicit chemical species (e.g., water). Moreover, atomic mass analysis has the shortcoming of not being applicable to large molecules because of the limitations of current annotation techniques. The second approach, Linear Programming (LP) analysis, avoids using model meta-data by checking for a weaker condition, stoichiometric inconsistency. But this approach suffers from false negatives and has large isolation sets (the set of reactions implicated in the stoichiometric inconsistency).

**Results:** We propose alternatives to both approaches. Our alternative to atomic mass analysis is moiety analysis. Moiety analysis uses model meta-data in the form of moieties present in chemical species. Moiety analysis avoids excessively detailed reaction specifications, and can be used with large molecules. Our alternative to LP analysis is Graphical Analysis of Mass Equivalence Sets (GAMES). GAMES has a slightly higher false negative rate than LP analysis, but it provides much better error isolation. In our studies of the BioModels Repository, the average size of isolation sets for LP analysis is 55.5; for GAMES, it is 5.4. We have created open source codes for moiety analysis and GAMES.

**Availability and Implementation:** Our project is hosted at https://github.com/ModelEngineering/ SBMLLint, which contains examples, documentation, source code files, and build scripts used to create `SBMLLint`. Our source code is licensed under the MIT open source license.

Supplementary information: None.

## 1 Introduction

Modeling is an essential part of science and engineering because of its ability to demonstrate complex phenomena and predict outcomes. In Systems Biology, many models are based on chemical reactions that specify how reactants are transformed into products. Herein, a reaction is specified by listing the chemical species (and their associated stoichiometry) for the reactants and the products. Such specifications are There are many similar public repositories of biological models such as central to kinetic models (e.g., Resat *et al*. [2009]) and flux balance analysis (e.g., Orth *et al*. [2010]).

The accuracy of reaction based models depends in large part on the correct specification of the reactions. Verifying reaction specifications has become quite challenging as reaction based models have grown in complexity. For example, BioModels (Le Novère *et al*. [2006]), a repository of literature-based physiologically and pharmaceutically relevant mechanistic models in standard formats, especially the Systems Biology Markup Language (SBML) (see (Hucka [2013]), contains over 600 curated models that range in size from tens to thousands of reactions.

MAMMOTh (Kazantsev *et al*. [2018]), CellML (Lloyd *et al*. [2008]), and BiGG (King *et al*. [2016]). The correctness of models in such public repositories is of particular concern since these repositories are often the starting point for new modeling projects.

How can we address the correctness of reaction based models as models grow in complexity? Our answer draws inspiration from approaches used in software engineering (Hellerstein *et al*. [2019]). If we view reaction based models as a kind of software, then the complexity of today’s reaction based models is comparable to the complexity of computer software in the early 1960s when programs were typically tens to a few thousand statements. Today, open source software such as Linux and the Apache Web Server have several million statements. This thousandfold increase in complexity is in part due to the development of sophisticated tools that automate error checking of software codes. One example is “linters” (e.g., Darwin [1988]) that check for errors such as a variable that is referenced before it is assigned and identifying unreachable code (e.g., a statement that follow a `return` statement). Both of these are examples of *static* error checking that is done by examining source codes without requiring them to be executed (Louridas [2006]).

Our goal is to develop linters that facilitate the development of complex reaction based models. A starting point is checking for mass balance errors. A mass balance error occurs if there is an incorrect specification of a reaction so that the total mass of the chemical species in the reactants differs from the total mass of the products. Checking for mass balance errors can be done directly on the model source without running simulations or any calculation of model outputs. Others have noted the importance of this kind of error checking (e.g., Medley *et al*. [2016], Clark *et al*. [2012], Rohr *et al*. [2010]).

We proceed with an example from (curated) BioModels. Considered is the model BIOMD0000000255, a model with 827 reactions. Fig. 1 displays 6 of the model’s 827 reactions using the syntax of the Antimony modeling language (Smith *et al*. [2009]). Reactions `v537` and `v601` are uni-uni reactions in which there is a single reactant and a single product, and all stoichiometries are one. For a uni-uni reaction to preserves mass balance, the mass of the reactant must equal the mass of the product. Thus, assuming mass balance, the masses of chemical species `c160, c154`, and `c86` are equal. However, reaction `v13` implies that the mass of `c154` is less than that of `c160` (since `c10` must have a non-zero mass). Hence, a mass balance error is present.

**Fig. 1.**
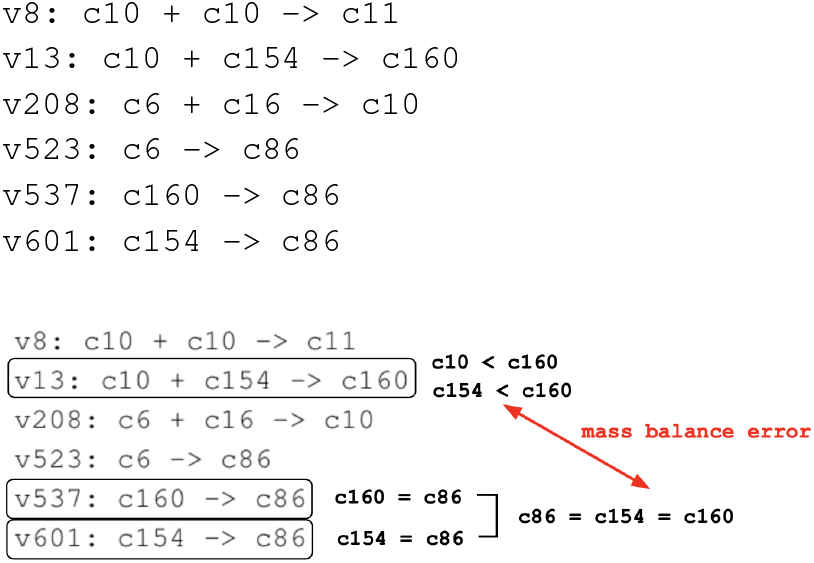
Snippet of BIOMD0000000255 that contains a mass balance error. Reactions are expressed using the syntax of the Antimony modeling language. Reactions are identified by the label that precedes the colon.

The foregoing analysis is tractable for 6 reactions. It would have been unmanageable for 827 reactions. This highlights the importance of **error isolation**, finding a small subset of reactions within a model that cause a mass balance error. While there has been substantial work on detecting mass balance errors, we are aware of only one prior effort that bears on error isolation, and this is done in a relatively indirect way.

Isolating mass balance errors is easy if error detection works at the granularity of individual reactions. Indeed, it would seem that if we know the chemical formula for each species in the reaction, then it should be trivial to check mass balance by comparing the total atomic mass of the reactants with the total atomic mass of the products. This approach is used by the `memote` system (Lieven *et al*. [2018]). Knowledge of formulas for chemical species is provided through **model meta data**, information about the elements of the biochemical model. Most commonly, this is done through annotations (e.g., Misirli *et al*. [2016], Neal *et al*. [2018]). We use the term **atomic mass analysis** to refer to approaches that use atom-level chemical formulas to detect mass balance errors.

There are two non-trivial challenges with atomic mass analysis. First, biochemical equations are not always written to provide balance at the level of atomic mass since many chemical species are assumed to be present in relatively constant concentrations (Nelson and Cox [2004]). We refer to such chemical species as **implicits**. For example, the hydrolysis of `ATP` is often written as `ATP → ADP`. Here, inorganic phosphate is implicit and so the equation does not account for one of the phosphate groups^1^ in the reactants. A more precise specification would be `ATP → ADP + Pi`, where `Pi` indicates a phosphate group. Still, this specification also fails mass balance because it does not account for small differences in atomic mass–the fact that the ***γ*** phosphate in `ATP` has a shared oxygen while the inorganic phosphate in the products does not. This can be addressed by including another implicit–water: ATP + H_2_0 → ADP + Pi (with appropriate accounting for the protons on the phosphate groups). Tools such as `memote` require that reactions are specified so that all chemical species are explicitly expressed in the reaction. Our experience is that such a requirement is quite burdensome for many modelers. One way to avoid this is to infer missing reactants (e.g., Swainston *et al*. [2011]), but this has the potential of inferring bogus reactions.

A second challenge posed by working in units of atomic mass is a limitation on what can be expressed in annotations. While annotations of small molecules are readily available, this is not the case for large molecules that are common in biochemistry. Examples include proteins, carbohydrate polymers, DNA, and RNA. A core challenge here is that large molecules can be in many chemical states (e.g., phosphorylation, protenation, and acetylation), and the state of a molecule must be considered in analyzing mass balance in units of atomic mass. Indeed, models of biological signaling consist mostly of reactions that describe changes in molecular states. Unfortunately, existing annotation systems do not have effective ways to describe such state changes.

The underlying issue in the foregoing is that we are working in units of atomic mass. It turns out that many mass balance errors can be detected without knowledge of the atomic formulas of chemical species. This is accomplished by detecting a slightly weaker condition–**stoichiometric inconsistency**, inconsistency in the relationships between reactions in a model. This approach is defined in terms of the stoichiometry matrix *N*, a matrix whose *ij*-th entry is the moles of a chemical species *i* which is produced by reaction *j*. This entry is negative if the reaction consumes species *i*. A stoichiometric inconsistency is present if there is no vector of positive masses *v* = {*v_i_*} such that *N^T^v* = 0.

The mass balance error in Fig. 1 is an example of a stoichiometric inconsistency in that for all positive masses assigned to `c154` and `c160`, there is no positive mass that can be assigned to `c10` that yields mass balance for reaction `v13`. Clearly, a stoichiometric inconsistency implies a mass balance error. However, there are mass balance errors that do not result in stoichiometric inconsistency. Although detecting stoichiometric inconsistency is a weaker condition than detecting mass imbalance, the former has great appeal because the detection of stoichiometric inconsistency *imposes no burden on the modeler*.

One way to detect stoichiometric inconsistency is to formulate a linear programming feasibility problem with the constraints *N^T^v* = 0 and *v* > 0 (Nikolaev *et al*. [2005], Gevorgyan *et al*. [2008]). We refer to this as `LP analysis`. Several tools implement this approach (e.g., Swainston *et al*. [2011], Schellenberger *et al*. [2011], Lund Steffensen *et al*. [2016]).

LP analysis does have some elements that relate to isolation of mass balance errors. Gevorgyan *et al*. [2008] propose the use of *minimal net stoichiometries* (the stoichiometries of chemical species implicated in a stoichiometric inconsistency) and *leakage modes* (linear combinations of reactions that result in stoichiometric inconsistencies). However, the resulting technical approach relies on mixed integer linear programming. This is computationally complex, and is only an indirect indication of the stoichiometric inconsistencies in reactions (although it does seem helpful for handling implicit molecules).

In summary, current approaches to detecting mass balance errors either use atomic mass analysis or LP analysis. Below we list issues with these approaches.

1. **I1: Model meta-data**. Atomic mass analysis requires that modelers annotate chemical species to provide atomic mass information.
2. **I2: Excessive detail**. Atomic mass analysis burdens modeler by requiring that reactions include implicit chemical species (e.g., protons, water, phosphate groups) so that mass balance can be calculated in units of atomic mass.
3. **I3: No large molecules**. Atomic mass analysis cannot address reactions with large molecules such as proteins, DNA, and RNA because of limitations of current annotation techniques (which do not address molecular states of large molecules).
4. **I4: Poor isolation**. LP analysis provides no practical way to isolate stoichiometric inconsistencies to individual reactions.
5. **I5: False negatives**. LP analysis detects stoichiometric inconsistencies and cannot detect all mass balance errors.

Issue I1 is inherent in any approach that requires model metadata, and I5 is inherent to any approach that focuses on stoichiometric inconsistencies. This paper proposes alternatives to mass balance analysis and LP analysis so as to address issues I2-I4.

We propose **moiety analysis**^2^ as an alternative to atomic mass analysis. Moiety analysis calculates mass in units of chemical groups or moieties instead of atomic mass. Because moieties can be defined in a flexible way that encompasses many chemically similar groupings of atoms, we avoid I2 and I3. Our alternative to LP analysis is **Graphical Analysis of Mass Equivalence Sets (GAMES)**. By using a combination of a graphical analysis and linear algebra, GAMES dramatically improves the identification of error causing reaction specifications (I4), although GAMES does have a slightly higher rate of false negatives than LP analysis. We have created open source implementations of moiety analysis and GAMES.

The remainder of this paper is organized as follows. Section 2 discusses moiety analysis and GAMES, and Section 3 evaluates these approaches using BioModels. Section 4 discusses the results and directions for future work.

## 2 Materials and methods

Our work is in part motivated by a need to improve on the error isolation provided by LP analysis. That is, once a mass balance error is detected, we want to find a small subset of reactions that caused the error. We refer to this subset as the **isolation set** for the mass balance error.

The isolation set provides a way to quantify the quality of error isolation. One metric is the cardinality of the isolation set, or the **isolation set size (ISS)**. Ideally, ISS is 1, which means that a single reaction has been identified as the cause of a mass balance error. In this case, we only need to examine the reactants and products of a single reaction (along with their stoichiometries). Atomic mass analysis produces an ISS of 1. As ISS increases, so does the difficulty of the analysis, because the relationships between chemical species shared by reactions become more complicated. For this reason, ISS quantifies the cognitive complexity for the resolution of mass balance errors.

We consider a second measure that normalizes ISS. This measure takes into account the fact that a model focuses on a part of a larger system. As such, there are often **boundary reactions** that create or destroy mass that transit the interface between what is modeled and what is not modeled.

The **normalized isolation set size (nISS)** is ISS divided by the number of non-boundary reactions. Clearly, nISS ranges from zero to one. nISS quantifies the effectiveness of an error isolation algorithm by indicating the fraction of the total number of reactions (less boundaries) that need to be considered.

This section describes two new approaches to isolating mass balance errors. Section 2.1 introduces moiety analysis, and Section 2.2 details Graphical Analysis of Mass Equivalence Sets (GAMES).

### 2.1 Moiety Analysis

Moiety analysis is an alternative to atomic mass analysis. Both require model meta-data, and so place a burden on the modeler to expose information. However, moiety analysis avoids issues with excessively detailed reactions (I2) and dealing with large molecules (I3).

Moiety analysis checks for mass balance in units of moieties instead of atomic mass. A moiety is one or more groupings of atoms that are chemically similar. For example, the three phosphate groups in ATP along with inorganic phosphate are similar chemically, although they have slightly different atomic formulas; so, we refer to all four as belonging to the phosphate moiety. In moiety analysis, the meaning of “chemically similar” is ultimately decided by the modeler, although there are well established guidelines in organic and biochemistry.

Moiety analysis examines the moiety structure of chemical species to ensure that there are equal counts of moieties in the reactants and products; implicit molecules are ignored in this analysis. To illustrate, consider the reaction `ATP → ADP + Pi`. Assume that there are two moieties: `A` is an adenosine moiety, and `Pi` is a phosphate moiety. We see that there is one `A` in both the reactants and the products, and there are three `Pis` in both the reactants and the products. So, moiety analysis detects no mass balance error.

Moiety analysis requires model meta-data that reveals the moiety counts of chemical species. This meta-data could be exposed through annotations. However, current annotation systems do not provide moiety information. Herein, we rely on a naming convention for chemical species to expose moiety structures.

Besides exposing moiety structures, we want a naming convention that is compatible with the SBML community standard (Hucka [2013]). This has two implications: (1) moiety names must follow the same rules as names of SBML chemical species, and (2) the only special character that can be used as a separator is an underscore (“_”).

Our naming convention uses single and double underscores as separators. This is best communicated by an example. Consider the moiety names `A` and `Pi`. Then the chemical species `ATP` is written as either `A__Pi__Pi__Pi` or `A_Pi_Pi_Pi`. Because it is common to have multiple instances of a moiety in a chemical species, we provide an alternative representation that includes a repetition count: `A__Pi_3`, where “_3” indicates that the phosphate is repeated three times, and a single underscore is used to separate the repetition count from the moiety name. When repetition counts are used in chemical species, then only a double underscore can be used to separate moieties.

The **occurrence count** of a moiety in reactants (or products) is the total count of the occurrences of the moiety in the chemical species of the reactants (or products) weighted by its stoichiometry. To illustrate, consider the first reaction of the Urea Cycle (Jackson [1986]) that has the reactants ammonia, bicarbonate, and two ATP molecules. We use the corresponding species names `Am, Bi`, and `A__Pi_3`. So, the reactant expression is `Am + Bi + 2 A__Pi_3`. The occurrence count of the moiety `A` is 2 because of the stoichiometry of `ATP` in the reactants. The occurrence count for `Pi` is 6 because there are three inorganic phosphates for each `ATP`.

We note in passing that, the atomic mass analysis is not sufficiently flexible to incorporate implicits as is done in moiety analysis. This is because such fine grain units do not allow for identification of chemical groups. One possible solution is to develop an algorithm that parses atomic structures for “implicit chemical groups” so that they could be removed from the analysis. However, there are subtleties in doing so since parsing must account for variants of chemical groups.

The flexibility provided by working in units of moieties has considerable benefits. In Section 1, we saw that analyzing `ATP → ADP + Pi` using atomic mass analysis requires the inclusion of the implicit water molecule in the reaction specification. Moiety analysis does not require the inclusion of such implicits, and so modelers can avoid unnecessarily detailed reaction specifications. Indeed, if `Pi` is specified as being implicit, the reaction can be further simplified to `A__Pi_3 → A__Pi_2`. Last, moiety analysis can be applied to large molecules since it is easy to represent changes in chemical state as changes in moiety structure. For example, if a protein is represented by the moiety `Prot`, then its phosphorylation is readily represented as `Prot_Pi`. This means that we can easily check mass balance in the phosphorylation reaction `Prot + A__Pi_3 → Prot__Pi + A__Pi_2`.

In summary, moiety analysis always produces an isolation set size (ISS) of one if there is a mass balance error. One caveat, however, is that moiety analysis imposes a burden on the modeler to expose model metadata that reveals the moiety structure of chemical species. In our approach, this meta-data is exposed through a naming convention. This seems to be a modest burden since the particulars of the names are at the discretion of the modeler.

### 2.2 Graphical Analysis of Mass Equivalent Sets (GAMES)

We can achieve excellent error isolation by requiring model meta-data (e.g., atomic mass analysis and moiety analysis). However, requiring meta-data imposes a burden on the modeler, and this requirement creates a barrier to analyzing existing models that have no meta-data. These concerns motivate analyzing models for stoichiometric inconsistencies, a slightly weaker condition than mass balance. Unfortunately, the existing approach to detecting stoichiometric inconsistencies, LP analysis (Nikolaev *et al*. [2005] and Gevorgyan *et al*. [2008]), has very poor error isolation in that all non-boundary reactions are in the isolation set.

Here, we introduce the Graphical Analysis of Mass Equivalence Sets (GAMES) algorithm for detecting stoichiometric inconsistencies. GAMES is designed to facilitate error isolation. Indeed, in our studies of BioModels, the average ISS of GAMES is less than 10% of the ISS of LP analysis.

The GAMES algorithm consists of three parts, as discussed in the next subsections. Section 2.2.1 describes the core algorithm, which we call basic GAMES or bGAMES. Section 2.2.2 describes xGAMES, which extends the ability of GAMES to detect mass balance errors by transforming the reaction network using linear algebra. Section 2.2.3 describes the GAMES approach to computing isolation sets.

#### 2.2.1 Detection With Basic GAMES (bGAMES)

bGAMES can be viewed as a kind of “inference engine” that attempts to construct inferences that contradict the assumption that all chemical species have positive mass. The inference steps rely on mass equality and inequality relationships implied by model reactions. If bGAMES infers that a chemical species does not have a positive mass, then the isolation set consists of the reactions used in the inference.

bGAMES uses insights such as those employed in the analysis of Fig. 1 to construct equality and inequality relationships between the masses of chemical species. For example in Fig. 1, `v537: c160 → c86` implies that the mass of `c160` must be the same as the mass of `c86`. This is an example of a **uni-uni** reaction, a reaction with one reactant and one product that have the same stoichiometries. A uni-uni reaction implies that the mass of the reactant equals the mass of the product.

The equality relationships implied by uni-uni reactions allow bGAMES to construct groupings of chemical species that have the same mass. We refer to such a grouping as a **Mass Equivalent Set (MEQ)**. In Fig. 1, reaction `v537` infers a MEQ that consists of the species `c160` and `c86`, which we denote by `{c86=c160}`. MEQs are expanded by transitivity. To illustrate, Fig. 1 also contains the MEQ `{c154=c86}` (implied by `v601`), and so by transitivity we have `{c86=c154=c160}`. Incorporating `v523` as well results in `{c6=c86=c154=c160}`. To account for all species in Fig. 1, we also have three singleton MEQs: `{c10}, {c16}` and `{c11}`.

bGAMES also infers mass inequalities. For example, two mass inequalities can be inferred from the reaction `v13: c10 + c154 → c160:` (a) the mass of `c160` is greater than the mass of `c10` and (b) `c160` is greater than `c154.v13` is an example of a **multi-uni** reaction, in which the reactants (or products) consist of two or more chemical species and there is a single chemical species as the product (or reactant).

The bGAMES inference engine uses uni-uni and multi-uni reactions to construct a directed graph that is then analyzed to detect stoichiometric inconsistencies. We refer to this as the **MEQGraph** since the nodes of the graph are MEQs. If (*X,Y*) is an arc in the MEQGraph, then the mass of MEQ *X* is less than the mass of MEQ *Y*. MEQs are constructed from the transitive closure of uni-uni reactions, and arcs are constructed from multi-uni reactions. Specifically, suppose there is a reaction `a + b → c`, with `a` in MEQ *X*, `b` in MEQ *Y*, and `c` in MEQ *Z*. Then, the MEQGraph has the arcs (*X,Z*) and (*Y,Z*).

Fig. 2 summarizes the bGAMES algorithm. bGAMES detects a stoichiometric inconsistency by finding a cycle in the MEQGraph. This is because a cycle implies a logical contradiction, that all MEQs in the cycle have a mass less than their own mass. It is straightforward for bGAMES to handle implicit chemical species. We simply add a pre-step that deletes implicits from reactions.

**Fig. 2.**
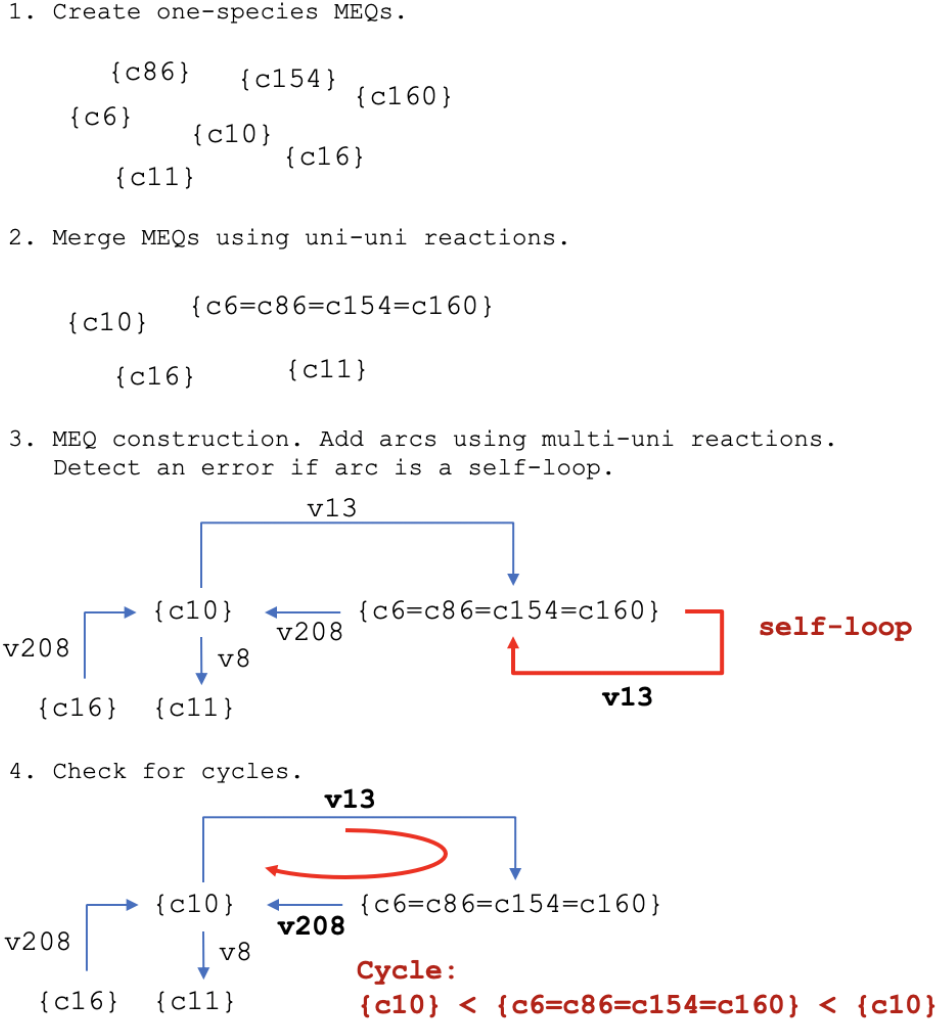
Steps in the bGAMES algorithm. The figure illustrates the data structures used in each step of bGAMES for the reactions in Fig.1. A mass balance error is detected if there is a loop in the graph since this implies a contradiction that some chemical species has a mass less than its own mass.

bGAMES does not detect *all* stoichiometric inconsistencies. This is because bGAMES does not analyze **multi-multi** reactions, reactions with more than one chemical species for both reactants and products. It turns out that despite this limitation, bGAMES still detects 75% of the models with stoichiometric inconsistencies that are detected by LP analysis in the BioModels repository. In large part, this is because many models contain few if any multi-multi reactions. For example, BIOMD0000000255 has 827 reactions, and no multi-multi reaction.

#### 2.2.2 Detection With Extended GAMES (xGAMES)

This section describes xGAMES, an extention of the bGAMES inference engine to detect and isolate stoichiometric inconsistencies that are not detected by bGAMES. The central idea of xGAMES is to perform certain transformations of reactions.

We illustrate this using BioModels BIOMD0000000167. (See Fig. 3.) Consider the following two reactions, `R2: Pstat_nuc → stat_nuc` and `R4: 2 Pstat_nuc → PstatDimer_nuc`. If these reactions are mass balanced, then the mass of `3 Pstat_nuc` must equal the sum of the masses of `stat_nuc` and `PstatDimer_nuc`. Put differently, we have mass balance for the hypothesized reaction `R2+R4: 3 Pstat_nuc → stat_nuc + PstatDimer_nuc`. We do not claim that this reaction is chemically feasible. Instead, we refer to this as a **pseudo reaction**. Even though it is not necessarily a real reaction, this pseudo reaction should be mass balanced. When mass balance holds, we use the term **mass balanced pseudo reaction**.

**Fig. 3.**
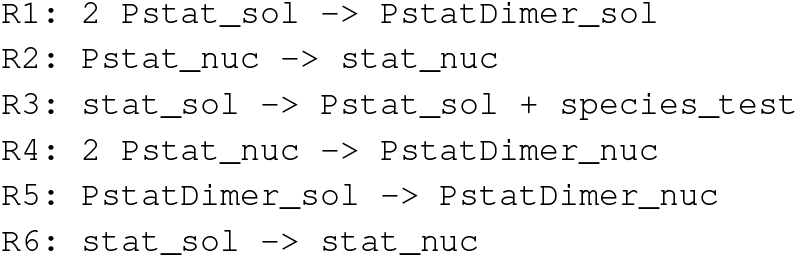
Reactions in BIOMD0000000167. The reaction names are changed to simplify the description. The complete model has two instances of `R6` with different kinetics.

Thus far, we consider just the summation of two reactions. We can generalize this. If a collection of reactions {*R_i_*} is mass balanced, then any linear combination of these reactions will be a mass balanced pseudo reaction.

The idea of reaction transformations is related to LP analysis. Recall that LP analysis detects a stoichiometric inconsistency if there is no positive vector of masses *v* such that *N^T^v* = 0, where *N* is the stoichiometry matrix (columns are reactions and rows are species). Such a *v* cannot be found if either: (a) the dimension of the column (species) null space of *N^T^* is 0; or (b) the column null space does not intersect the subspace where *v* is positive. xGAMES uses matrix decomposition to test for these conditions and do error isolation by reporting the reactions that result in a mass balance error.

xGAMES works with a new stoichiometry matrix (*N*) whose rows are MEQs instead of chemical species. Doing so allows us to eliminate uni-uni reactions since this information is already encoded in the MEQs. This is illustrated in Tab. 1. The columns are the pseduo reactions `PR3, PR1`, and `PR4` that correspond to the original reactions `R3, R1`, and `R4`. For example `PR1: 2{Pstat_sol} → {PstatDimer_sol=PsatDimer_nuc}`. Note that every column contains at least one negative value and at least one positive value. This is because every reaction has at least one reactant and at least one product.

**Table 1.** Stoichiometry matrix for pseudo reactions. Rows are MEQs. Columns are pseudo reactions that are numbered corresponding to the reactions in Fig. 3. Cells are the stoichiometry of a MEQ in the products minus the stoichiometry of the MEQ in the reactions. Uni-uni reactions are not included since they are used to construct the MEQs.

| MEQ | PR3 | PR1 | PR4 |
| --- | --- | --- | --- |
| {species_test} | 1 | 0 | 0 |
| {Pstat_sol} | 1 | -2 | 0 |
| {PstatDimer_sol=PstatDimer_nuc} | 0 | 1 | 1 |
| {Pstat_nuc=stat_nuc=stat_sol} | -1 | 0 | -2 |

Next, xGAMES transforms *N* into reduced column echelon form using standard techniques from linear algebra such as LU decomposition (Horn and Johnson [1985]). We denote this transformed matrix by 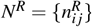. Tab. 2 displays *N^R^* for our running example. We can see that the leading entry in each column is one and 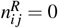 for *j* > *i*, as required by reduced column echelon form. Also, as with Tab. 1, the columns of Tab. 2 are pseduo reactions. Specifically, the columns of *N^R^* are linear combinations of the columns of *N*. For example, 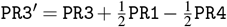.

**Table 2.** Reduced column echelon form for the running example. The prime symbol indicates a reaction formed by linear combinations of these in Tab. 1.

| MEQ | PR3' | PR1' | PR4 |
| --- | --- | --- | --- |
| {species_test} | 1 | 0 | 0 |
| {Pstat_sol} | 0 | 1 | 0 |
| {PstatDimer_sol=PstatDimer_nuc} | 0 | 0 | 1 |
| {Pstat_nuc=stat_nuc=stat_sol} | 0 | -1 | -2 |

xGAMES detects a mass balance error using the following decision criteria:

- **Detection Criteria (DC):** A mass balance error is present if there is a linear combination of columns of *N* whose non-zero values all have the same sign.

We note in passing that DC relates to the concept of a leakage mode in Gevorgyan *et al*. [2008], which is a linear combination of reactions that results in an empty set of reactants or products.

DC detects mass balance errors by identifying reactions for which mass is either created or destroyed. A column of the stoichiometry matrix in which all non-zero values are positive describes a reaction that has products and no reactants. That is, mass is created. Analogously, a column in which all non-zero values are negative describes a reaction that has reactants and no products, and so mass is destroyed.

xGAMES detects a stoichiometric inconsistency by inferring a logical contradiction. The inference engine starts by assuming that the original reactions are mass balanced. From the foregoing, we know that linear combinations of mass balanced reactions result in a mass balanced pseudo reactions. So, if some linear combination of reactions in the model results in DC being satisfied, then we know that the original set of reactions is *not* mass balanced.

Fig. 4 depicts the steps in the xGAMES algorithm. It stops when a mass balance error is detected. This is because it is computationally expensive to report all mass balance errors, and likely very confusing to the modeler as well since one incorrect reaction specification may cause multiple mass balance errors.

**Fig. 4.**
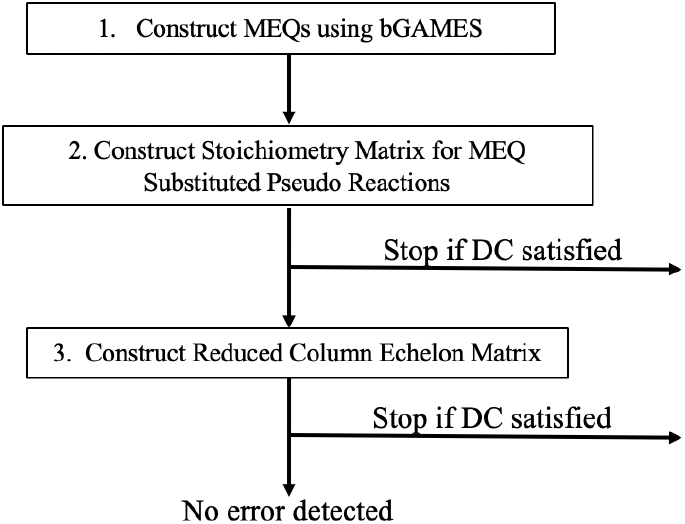
Steps in the xGAMES algorithm. The algorithm stops when the first error is encountered using the decision criteria (DC).

xGAMES occasionally fails to detect a stoichiometric inconsistency when one exists; that is, xGAMES can produce more false negatives than LP analysis. To understand why, recall that a stoichiometric inconsistency is present if (1) there is no vector of masses *v* such that *N^T^v* = 0 (where *N* is the stoichiometry matrix) such that (2) *v* > 0. LP analysis checks precisely for these two conditions. The xGAMES decision criteria DC detects violations of condition (1) if *N^T^* has a trivial null space. However, DC only approximates detection of violations to condition (2). Specifically, DC looks for columns where the non-zero values have the same sign. This is a necessary but not a sufficient condition for detecting violations of *v* > 0.

Despite this limitation, our experience with applying xGAMES to BioModels has been excellent–the xGAMES false negative rate with respect to LP analysis is under 5%.

#### 2.2.3 Computing GAMES Isolation Sets

This section describes how GAMES calculates isolation sets, the set of reactions that relate to a stoichiometric inconsistency. The approach taken depends on whether the mass balance error is detected by bGAMES or xGAMES.

If a mass balance error is detected by bGAMES, then there is a cycle in the MEQGraph. The isolation set consists of the reactions that are either: (a) multi-uni reactions that construct an arc in the cycle or (b) uni-uni reactions that construct a MEQ in the cycle. For example, in Fig. 2, there is a cycle with arcs labelled with reactions `v13` and `v208`; the MEQs are `{c10}` and `{c6=c86=c154=c160}`. The latter MEQ is constructed from the uni-uni reactions `v523, v537`, and `v601`. Thus, the isolation set is `{v13, v208, v523, v537, v601}`.

A different kind of calculation is required to determine the isolation set for an error detected by xGAMES. Recall that the xGAMES detection criteria (DC) is that there is a column in the stoichiometry matrix such that all non-zero values are of the same sign (i.e., mass has either been created or lost). The isolation set is the reactions in the linear combinations that result in such a column.

How do we recover these reactions? In Step 2, the stoichiometry matrix is constructed from the MEQ substituted pseudo reactions. So, the isolation set for a column that satisfies DC is the original reaction for the same column plus the uni-uni reactions for the MEQs in the pseudo reaction.

The matrix in Step 3 is the reduced column echelon form of the stoichiometry matrix in Step 2. The analysis here is more involved. Let *N* be the stoichiometry matrix constructed in Step 2. We use LU decomposition (Horn and Johnson [1985]) to factor *N^T^* into *PLU*, where *P* is a permutation matrix and *L, U* are lower and upper triangular matrices, respectively. With rare exception *L* is invertible. (The exceptions have been related to numerical issues similar to those encountered with LP analysis.) So, *U* = *L*^−1^ *P*^−1^ *N^T^*, and *U^T^* = *NP*(*L*^−1^)^*T*^. The columns of *P*(*L*^−1^)^*T*^ reveal the linear combinations of reactions of *N* that result in the columns echelon form. This is readily extended to reduced column echelon form by elementary matrix operations so that we have a matrix that informs us of the linear combinations of columns of *N* that are used to construct the reduced column echelon matrix *R*. The isolation set is the set of reactions that correspond to the columns in this linear combination that have non-zero multipliers.

## 3 Results

This section evaluates moiety analysis and GAMES in the context of the curated SBML models in the BioModels repository. For moiety analysis, the focus is on the ease with which existing models can be transformed to use the moiety analysis naming convention. For GAMES, we address runtimes, false negatives compared with LP analysis, and the size of isolation sets. Our evaluations are done using `SBMLLint`.

The evaluations done in this section make use of the 651 curated SBML models in BioModels. We focus on curated models because of the human effort expended to validate their correct operation. These models address a wide range of biological processes, including metabolism, signaling, and motility. We have applied LP analysis to the 651 curated models, and found that approximate 20% of the models have at least one stoichiometric inconsistency. While this does not necessarily mean that the models are in error in terms of the studies for which they were designed, it does raise concerns for researchers considering reuse of these models for new studies.

Tab. 3 displays statistics for the types of reactions of these models. We see that multi-multi reactions are relatively uncommon, accounting for about 10% of all models. If more multi-multi reactions were present, it is likely that bGAMES would have a higher rate of false negatives since bGAMES does not consider multi-multi reactions.

**Table 3.** Average number (± standard error) of reactions by type in the 651 curated reactions from BioModels. Count is the average number of each reaction type, and Fraction is the average fraction of each reaction type across models.

| Measure | Uni-Uni | Multi-Uni | Multi-Multi | Boundary | Total |
| --- | --- | --- | --- | --- | --- |
| Count | 12.1±1.8 | 13.8±2.1 | 4.9±1.0 | 5.8±0.5 | 36.7±3.6 |
| Fraction | 0.36 | 0.25 | 0.10 | 0.29 | 1.00 |

### 3.1 SBMLLint

We have created an open source implementation of moiety analysis and GAMES algorithm, which is available in the `github` repository https://github.com/ModelEngineering/SBMLLint. The term “lint” in the name comes from software engineering, and refers to tools that do static error checking. Unlike systems such as `memote` that do static error checking as part of a bigger system, `SBMLLint` operates as a stand-alone package that can be used in isolation. `SBMLLint` takes as input an SBML XML file or a model in the user-friendly antimony modeling language (Smith *et al*. [2009]). `SBMLLint` can be run at the command line or in a python environment, such as a Jupyter Notebook (Pérez and Granger [2007]).

### 3.2 Evaluation of Moiety Analysis

We have found that a non-trivial fraction (15%-20%) of the curated models in BioModels use names of chemical species that are nearly compliant with the moiety analysis naming convention. Consider the model in Fig. 3 that has the species names `Pstat_sol, PstatDimer_sol, stat_sol, Pstat_nuc, PstatDimer_nuc`. These names can be deconstructed into string expressions so that the names are structured as follows: (a) “`Pi`” or “” followed by (b) `stat` followed by (c) `Dimer` or “” followed by (d) `_sol` or `_nuc`. In essence, the model in Fig. 3 uses a different naming convention from the one we propose, but species names still have a moiety structure (although `sol` and `nuc` seem to refer to compartments, not chemical constituents). There are many similar situations in the 651 curated models.

The supplemental material details two case studies of converting existing models in BioModels to using moiety analysis.

### 3.3 Evaluation of GAMES

This section provides insights into the effectiveness of the GAMES algorithm. The supplemental material details two case studies of GAMES analysis done on models in BioModels.

To provide broader insights into GAMES, we conducted an analysis of the 651 curated models, applying both LP analysis and GAMES to these models. Tab. 4 displays the results, including separate statistics for bGAMES and xGAMES. We see that both bGAMES and xGAMES have a longer runtime than LP analysis. Further optimization of the GAMES code will likely reduce the time for a GAMES analysis. That said, these optimizations may be unnecessary, at least with current model complexities, since the average time per model is a few hundred milliseconds, which is quite acceptable for use in an interactive tool.

**Table 4.** Comparison of three approaches to detect mass balance errors for 651 curated models in BioModels. The first two data columns report the time to run the algorithms for all models and per model, respectively. Detections is the number of models found to have a stoichiometric inconsistency. False negatives are reported with respect to LP analysis.

| Algorithm | Total sec | per Model sec | Detections | False Negatives |
| --- | --- | --- | --- | --- |
| LP analysis | 66.9 | 0.10 | 143 | reference |
| bGAMES | 123.2 | 0.19 | 109 | 23.8% |
| xGAMES | 220.5 | 0.34 | 136 | 4.9% |

Next, we study the error isolation. We consider two measures. The first is isolation set size (ISS), the number of reactions associated with the detection of a stoichiometric inconsistency. Ideally, ISS is 1 so that error isolation points to a single reaction that causes the mass balance error. As the ISS grows, so does the difficulty of error remediation. A second measure of interest is the normalized size of the isolation set (nISS). nISS is ISS divided by the number of non-boundary reactions, a quantification of the extent to which the algorithm simplifies error isolation.

Tab. 5 displays ISS and nISS for LP, bGAMES, and xGAMES. We see that the LP analysis results in an average ISS of 55.5 reactions. It is doubtful that the underlying cause of a stoichiometric inconsistency can be determined with an ISS of 55.5, and so remediation would be nearly impossible. In contrast, ISS for bGAMES is approximately 4 and for xGAMES it is a bit more than 5.

**Table 5.**
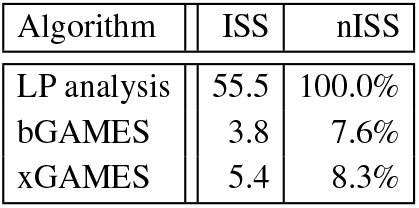
Isolation effectiveness for LP, bGAMES, and xGAMES. bGAMES and xGAMES reduce the size of isolation sets by a factor of more than 10 compared with LP analysis.

In summary, LP analysis provides excellent detection of stoichiometric inconsistencies, but very poor error isolation. Indeed, in our analysis of BioModels, the modeler would have to manually review an average of 55 reactions to resolve a stoichiometric inconsistency. In contrast, GAMES produces an isolation set of approximately 5 reactions instead of 55 (although GAMES has a a modest 4% rate of false negatives compared with LP analysis). The execution time to run GAMES is longer than that for LP analysis, but the code is still being optimized. And, even with the unoptimized codes, GAMES typically has sub-second execution times for BioModels.

## 4 Discussion

The growing complexity of reaction-based models makes it challenging to detect and resolve model errors such as mass balance errors in reaction specifications.

Tab. 6 summarizes how approaches to detecting mass balance errors fare relative to issues I1-I5. Most fundamentally, there is a divide between approaches that require model meta-data and those that do not. Providing model meta-model places a burden on the modeler to expose the meta-data (I1). However, by exposing meta-data, false negatives can be avoided (I5). Requiring atomic mass meta-data, as in atomic mass analysis, imposes additional burdens: excessively detailed reaction specifications (I2) and an inability to analyze reactions with large molecules (I3). Moiety analysis avoids I2 and I3 by using a different kind of meta-data – the moiety composition of chemical species.

**Table 6.** Comparison of approaches to mass balance errors. An “x” indicates that the issue is not addressed;a check indicates that the issue is addressed.

| Issue addressed | Atomic mass analysis | LP analysis | Moiety analysis | GAMES |
| --- | --- | --- | --- | --- |
| I1: Require meta-data | ✗ | ✓ | ✗ | ✓ |
| I2: Excessive detail | ✗ | ✓ | ✓ | ✓ |
| I3: No large molecules | ✗ | ✓ | ✓ | ✓ |
| I4: Poor isolation | ✓ | ✗ | ✓ | ✓ |
| I5: False negatives | ✓ | ✗ | ✓ | ✗ |

Techniques that do not require model meta-data do not check for mass balance errors per se; rather, they detect stoichiometric inconsistencies. This is a weaker condition than the mass balance, thus the presence of false negatives (I5) is inherent in these approaches. Beyond this, a major shortcoming of existing approaches to stoichiometric inconsistencies is the absence of effective techniques for problem isolation (I4). In our studies of BioModels, LP analysis has an average isolation set size (ISS) of 55.5 reactions. Such a large isolation set makes problem resolution extremely difficult. In contrast, GAMES has an average ISS of 5.4 reactions. The rate of false negatives for GAMES is slightly higher than for LP analysis because GAMES has a weaker test for stoichiometric inconsistency. This situation can be addressed by using LP analysis for error detection, and GAMES for error isolation.

Our interest in static error checking of reaction-based models naturally leads to considerations of charge balance. This can be approached using annotations (e.g., `memote`). Doing so has the same concerns as with mass balance – demanding excessively detailed reaction specifications (e.g., no “stray” protons), the burden of requiring model meta-data, and the limitations on what can be annotated. An alternative is to use moiety analysis. Specifically, if moieties are defined so that all atomic variants have the same charge, then moiety balance implies charge balance. Unfortunately, we know of no counterpart to stoichiometric inconsistency for charge balance because there is no constraint on charges akin to the constraint that all chemical species have a positive mass.

We are pursuing a number of directions for future work. For moiety analysis, we are exploring how the structure of names can be used to expose model meta-data other than the moieties of a chemical species, such as compartments and molecular states (e.g., misfolded proteins). For GAMES, we are investigating ways to reduce both runtimes and the rate of false negatives relative to LP analysis. More broadly, we are interested in providing modelers with insights into preferred practices or “model patterns,” which we might be able to infer from analyzing many models.

## Acknowledgements

We greatly appreciate the technical assistance provided by Valentina Staneva for her help with python linear algebra packages and related mathematical techniques. We are also indebted to Herbert M. Sauro for his thoughtful comments on earlier drafts.

## Footnotes

1 In this analysis, it is important to distinguish a phosphate group, which may be bound to one or more carbons, from an inorganic phosphate that does not have a carbon bond.

2 We use the term “moiety” as a generic reference to a chemical group.

