## Supplemental Material for "Detecting and Isolating Mass Balance Errors in Reaction Based Models in Systems Biology"

#### 1 Details of Moiety Analysis

Fig. S1 displays the moiety analysis algorithm for detecting mass balance errors. The first two steps compute occurrence counts for each moiety in the reactants and products. A mass balance error is detected if the occurrence count of a moiety in the reactants differs from that in the products. This check does not include moieties that the modeler specifies as implicit. For example, if the modeler indicates that `Pi` is implicit, then moiety analysis will *not* detect a mass balance error in the reaction `A__Pi_3` $\rightarrow$ `A__Pi_2`. By so doing, modelers can further simplify reaction specifications.

```
reactants[moiety] = reactant occurrence count of moiety
products[moiety] = product occurrence count of moiety
for unique moiety in reactants and products
  if moiety is not implicit
    if reactants[moiety] != products[moieties]
      report error
```

Fig. S1: *The moiety analysis algorithm for detecting mass imbalance for a reaction. Reactants and products are lists of chemical species with names structured by the moieties they contain. A reaction has a mass balance error if there is a moiety whose occurrence count in the products differs from that in the reactants.*

#### 2 Details of GAMES

The following provides more details about the steps in bGAMES.

1. Initialization is done so that there is a MEQ for each chemical species.

2. MEQs are merged. This is done using uni-uni reactions to discover mass equivalences and detecting non-null intersections between sets. By merging MEQs A and B, we mean that a new MEQ C is formed such that $C = A \cup B$, and $A, B$ are deleted.

3. Arcs are added between nodes (MEQs) based on multi-uni reactions to indicate strict inequality of masses.

4. The MEQGraph is checked for cycles. A cycle in the MEQGraph implies a logical contradiction, that some chemical species has a mass less than its own mass.

This is accomplished as follows: (a) delete from the MEQGraph all nodes (and their incident arcs) that are singleton implicits and (b) delete implicits from the remaining nodes and any incident arc for that implicit. For example, if `c6` in reaction `v208` is an implicit, then we delete the arcs labelled `v208` and change the node `{c6=c160=c86=c154}` to `{c160=c86=c154}`. This leaves the self-cycle caused by `v13`, and so a mass balance error is still detected. Its isolation set consists of `v13` and the uni-uni reactions that construct the MEQ `{c160=c86=c154}`.

There are a few caveats to the bGAMES algorithm. First, as with the LP Algorithm, bGAMES only addresses stoichiometric inconsistencies, not the full range of possible mass balance errors. To the best of our knowledge, the only way to expand beyond stoichiometric inconsistencies is for the modeler to provide more information, such as is done with moiety analysis.

### 3 Moiety Analysis Case Studies

We proceed with case studies of converting two curated models to use the moiety analysis naming convention. Considered first is BIOMD0000000140 (Hoffmann *et al.* [2007]), a model of temporal control and selective gene activation in mammalian cells. The model has 45 reactions and 65 parameters and chemical species. Many of the chemical species are already named in the manner required by moiety analysis; even better, many of the reactions are moiety balanced. For example, `v1:  NFkB + IkBalpha -> IkBalpha_NFkB`. However, a first attempt at applying `SBMLLint` using moiety analysis results in 23 non-boundary reactions with mass balance errors. Sixteen of the errors result from four implicits: `IkBalpha`, `IkBbeta`, `IkBeps`, and `IKK`. In addition, there are seven reactions that use the name of the chemical species to indicate its compartment. An example is transport into the nucleus, such as `IkBbeta -> IkBbeta_nuc`. Fortunately, this can be handled by treating the location (e.g., `nuc`) as an implicit. Once we specify the aforementioned implicits to `SBMLLit`, there is no mass balance error.

A second case study is BIOMD0000000293 (Proctor *et al.* [2010]), which models the ubiquitin-proteasome system and its role in protein aggregation and degrading damaged proteins. The model has 316 reactions, including 10 boundary reactions. As with the previous example, a significant number of reactions (about 50) already use the moiety analysis naming convention and are mass-balanced. An example is `Parkin_asyn_dam_Ub6 + Proteasome -> asyn_dam_Ub6_Proteasome + Parkin`. However, in many cases modest edits are required to expose the underlying structure of names. Examples include: renaming `ATP` as `A__Pi_3`, `NatP` (native proteins) as `Nat__Prot`, and

Fig. S2: *Example of GAMES report when running* `SBMLLint`*. The report has four sections. The first section displays the isolation set. The remaining sections provide details of how the isolation set was constructed.*

```
We detected a mass imbalance
:  -> species_test

from the following isolation set.

1. statPhosphorylation: stat_sol -> Pstat_sol + species_test
2. PstatDimerisation: 2.00 Pstat_sol -> PstatDimer_sol
3. PstatDimerisationNuc: 2.00 Pstat_nuc -> PstatDimer_nuc

----------------------------------------------------------------------
----------------------------------------------------------------------

These uni-uni reactions created mass-equivalence.
(The chemical species within a curly bracket have the same atomic mass.)

{Pstat_nuc=stat_nuc=stat_sol} is inferred by:
4. stat_export: stat_sol -> stat_nuc
5. statDephosphorylation: Pstat_nuc -> stat_nuc

{PstatDimer_nuc=PstatDimer_sol} is inferred by:
6. PstatDimer__import: PstatDimer_sol -> PstatDimer_nuc

----------------------------------------------------------------------

Based on the uni-uni reactions above, we create
mass-equivalent pseudo reactions.
(pseudo 1.) statPhosphorylation:
           {Pstat_nuc=stat_nuc=stat_sol} -> {Pstat_sol} + {species_test}
(pseudo 2.) PstatDimerisation: 2.00
                          {Pstat_sol} -> {PstatDimer_nuc=PstatDimer_sol}
(pseudo 3.) PstatDimerisationNuc: 2.00
        {Pstat_nuc=stat_nuc=stat_sol} -> {PstatDimer_nuc=PstatDimer_sol}

----------------------------------------------------------------------

An operation between the pseudo reactions:
1.00 * statPhosphorylation + 0.50 * PstatDimerisation -
0.50 * PstatDimerisationNuc

will result in empty reactant with zero mass:

:  -> {species_test}
```

`UCHL1damaged` as `UCLH1__damaged`. We see that strings such as `Nat` and `damaged` describe the molecule structure, not its mass. We run `SBMLLint` with the following implicits: `Agg`, `dam`, `damaged`, `Mis`, `misfolded`, `Nat`, `P`, `Proteasome`, `ROS`, `Seq`, `Source`, and `upreg`. This results in 146 reactions with a mass balance error. A common themes in these reactions is not properly expressing the mass present in a molecule aggregation, disaggregation, or degredation. For example, the original model has the reaction `AggP2 -> AggP1 + MisP`, which we translate into `Agg__Prot_2 -> Agg__Prot + Mis__Prot`. This is mass balanced.

However, `AggP1 -> 2.00 MisP`, which we translate to `Agg__Prot_1 -> 2.00 Mis_Prot` is not mass balanced since there are two `Prot` in the products but only one in the reactants.

The foregoing case studies suggest that there is a modest burden imposed on modelers to abide by the moiety analysis naming convention.

### 4 GAMES Case Studies

Our first case study is the model in Fig. 3, BIOMD000000167 (Li *et al.* [2010]). The full model has seven reactions and seven chemical species. This model has a mass imbalance, a fact that is far from obvious. Even more surprising is the difficulty of finding the cause of the mass imbalance by a manual inspection.

Fig. S2 displays the report that GAMES produces when `SBMLLint` analyzes BIOMD000000167. The report has four sections. The first section reports the reactions in the isolation set. The remaining three sections detail how the isolation set is constructed. Section two lists the uni-uni reactions from which MEQs are constructed, and section three displays the MEQ substituted pseudo reactions. Finally, section four displays a linear combination of the pseudo reactions in section three that result in a mass balance error. In this case, the mass balance error is detected as the creation of mass. Our suspicion is that the error relates to the reaction `statPhosphorylation` since it seems curious that `species_test` *and* `Pstat_sol` are created from `stat_sol`.

Our second case study is a much larger model, BIOMD000000049 (Sasagawa *et al.* [2005]). This model simulates extracellular-signal-regulated kinase (ERK) signalling networks. The model has 150 reactions and almost 100 chemical species. Here, the isolation set consists of five reactions: `J44`, `J46`, `J88`, `J112`, and `J164`. The GAMES report shows that from these reactions we can infer the pseduo reaction `proteosome -> `.
